# Place cells are non-randomly clustered by field location in CA1 hippocampus

**DOI:** 10.1101/2022.03.15.484476

**Authors:** Hannah S. Wirtshafter, John F. Disterhoft

## Abstract

A challenge in both modern and historic neuroscience has been achieving an understanding of neuron circuits, and determining the computational and organizational principles that underlie these circuits. Deeper understanding of the organization of brain circuits and cell types, including in the hippocampus, is required for advances in behavioral and cognitive neuroscience, as well as for understanding principles governing brain development and evolution. In this manuscript, we pioneer a new method to analyze the spatial clustering of active neurons in the hippocampus. We use calcium imaging and a rewarded navigation task to record from 100s of place cells in the CA1 of freely moving rats. We then use statistical techniques developed for and in widespread use in geographic mapping studies, global Moran’s I and local Moran’s I to demonstrate that cells that code for similar spatial locations tend to form small spatial clusters. We present evidence that this clustering is not the result of artifacts from calcium imaging, and show that these clusters are primarily formed by cells that have place field around previously rewarded locations. We go on to show that, although cells with similar place fields tend to form clusters, there is no obvious topographic mapping of environmental location onto the hippocampus, such as seen in the visual cortex. Insights into hippocampal organization, as in this study, can elucidate mechanisms underlying motivational behaviors, spatial navigation, and memory formation.

## Introduction

The brain’s organization has long been one of the questions driving neuroscience research. In the past 100 years, recording and imaging techniques have allowed the examination of increasingly small cellular circuits and cell clusters (Aharoni and Hoogland, 2019a; Boyden, 2015; Gray et al., 1995; Jones, 2007; Scott et al., 2018; Verkhratsky and Parpura, 2014). These discoveries have led to important insights on, among other things, functional connectivity (Dombeck et al., 2009; Robert et al., 2021; Shin et al., 2020; Stratford and Wirtshafter, 2012; Wirtshafter and Wilson, 2019), plasticity and learning (Cai et al., 2016; Josselyn and Tonegawa, 2020; Robert *et al*., 2021; Weible et al., 2003), and brain development and evolution (Muessig et al., 2015; O’Reilly et al., 2015; Tosches, 2017). Although the hippocampus (HPC) receives topographically ordered inputs (Raisman et al., 1966; Risold and Swanson, 1996; Swanson and Cowan, 1977) and outputs to other regions (e.g. the lateral septum) (Ferreira-Fernandes et al., 2019; Linke et al., 1995; Swanson, 1977; Wirtshafter and Wilson, 2021) in an ordered fashion, the fine scale organization of the HPC—if it exists—remains to be elucidated.

If “structure determines function” and the ability of a neural circuit to perform a computation or function derives from its organizational principles, it becomes even more essential to understand these principles in order to understand the functions of the brain. Organizational principles guiding the organization and topography of many brain regions, including the motor cortex (Dombeck *et al*., 2009), visual cortex (Swindale, 1996; Van Essen and Maunsell, 1983), and several subcortical regions (Nakano, 2000; Stratford and Wirtshafter, 1990; Wirtshafter and Sheppard, 2001) have been established, however, there has been little consensus regarding if and how place cells in the hippocampus are organized (Dombeck et al., 2010; Eichenbaum et al., 1989; Franca and Monserrat, 2019; Hampson et al., 1999; Muller et al., 1987; Nakamura et al., 2010; O’Keefe et al., 1998; Pavlides et al., 2019; Redish et al., 2001).

The presence of micro-level organization in the hippocampus has the potential to provide insight on the circuit and systems level processes that govern memory formation, associative conditioning processes, and spatial navigation (Eichenbaum et al., 1992; Hasselmo, 2015; McNaughton et al., 2006; Moore et al., 2021; Olton et al., 1979; Sweis et al., 2021). One method to investigate cellular organization of the HPC is to examine whether place cell firing fields have any topographic organization. Pyramidal cells in the hippocampus, which increase their firing rates at specific locations during navigation (Knierim, 2015; O’Keefe, 1976; O’Keefe and Nadel, 1978), are believed to be involved in spatial learning (McNaughton *et al*., 2006; O’Keefe and Nadel, 1978; Wikenheiser and Redish, 2015a), memory encoding and recall (Farovik et al., 2010; Jadhav et al., 2012; O’Neill et al., 2010; Poucet et al., 2000; Wilson and McNaughton, 1994), and reinforcement and reward seeking (Gauthier and Tank, 2018; McGlinchey and Aston-Jones, 2018; Michon et al., 2019; Wikenheiser and Redish, 2015b; Wirtshafter and Wilson, 2020). Additionally, neurons that act as place cells in one environment also have the capacity to encode other parameters (such as time (Eichenbaum, 2014; Mau et al., 2018), conditioned and unconditioned stimuli and reward (McEchron and Disterhoft, 1997; Weible *et al*., 2003; Wirtshafter and Wilson, 2019), and, in humans, even abstract concepts (Bausch et al., 2021; Quiroga, 2012; Suthana et al., 2015)) in different environments (Schimanski et al., 2013; Shan et al., 2016). Determining the spatial organization of place cells in the hippocampus may therefore also elucidate the cellular circuits underlying many types of processing and cognition.

Previous methods of cellular recording, such as electrophysiology, do not allow the exact spatial organization of cells to be determined or analyzed. With the increasing use of calcium imaging techniques using viral GCaMPs, the relative positions of individual cells can be studied (Aharoni et al., 2019; Silva, 2017). However, using calcium imaging in mice yields a low number of place cells (a previous study examining CA1 organization recorded an average of 17 place cells per session (Dombeck *et al*., 2010)), which prevents a thorough analysis of place cell firing patterns with respect to their relative locations in CA1. The expansion of hippocampal CA1 calcium imaging into rats has allowed the imaging of a larger number of total cells per session, with almost 80% of imaged cells displaying place fields (Wirtshafter and Disterhoft, 2022). The ability to image 100s of place cells at once, and to analyze their corresponding place fields, provides an unprecedented opportunity to examine the spatial organization of individually imaged CA1 place cells.

In this manuscript, we pioneer a new method to analyze the spatial clustering of active neurons in the hippocampus. We use calcium imaging and a rewarded navigation task to record from 100s of place cells in the CA1 of freely moving rats. We then use statistical techniques developed for and in widespread use in geographic mapping studies, global Moran’s I and local Moran’s I (Anselin, 1995; Bivand and Wong, 2018; Cliff and Ord, 1981; Moran, 1950; Rushton, 2003), to demonstrate that cells that code for similar spatial locations tend to form small spatial clusters. We present evidence that this clustering is not the result of artifacts from calcium imaging, and show that these clusters are primarily formed by cells that have place field around previously rewarded locations. We go on to show that, although cells with similar place fields tend to form clusters, at the scale examined in this study, there is no obvious topographic mapping of environmental location onto the hippocampus, such as seen in the visual cortex.

## Results

Four male Fischer 344 x Brown Norway rats were injected with AAV9-Syn- GCaMP7c below the CA1 cell layer, and a 2mm grin lens was implanted into the CA1. After recovery and confirmation of visible cells, animals were then food restricted to 85% body weight and trained on a 2.64m linear track. Animals were rewarded with grain pellets on alternating sides of the linear track and quickly learned to run back and forth on the track (figure 1A). A visit to one reward well signified a trial. Animals were run for a minimum of 30 minutes or 50 trials per day, whichever came later, with total track time not exceeding one hour per session. Six sessions were analyzed from each animal (see methods), with a total of 5761 neurons recorded over the 24 sessions, with a maximum of 428 cells from a single session (mean of 239.6+-90.0 cells). For each recording, we optimized the number of cells being imaged rather than the maintenance of the same field of view or focal plane.

**Figure 1:**
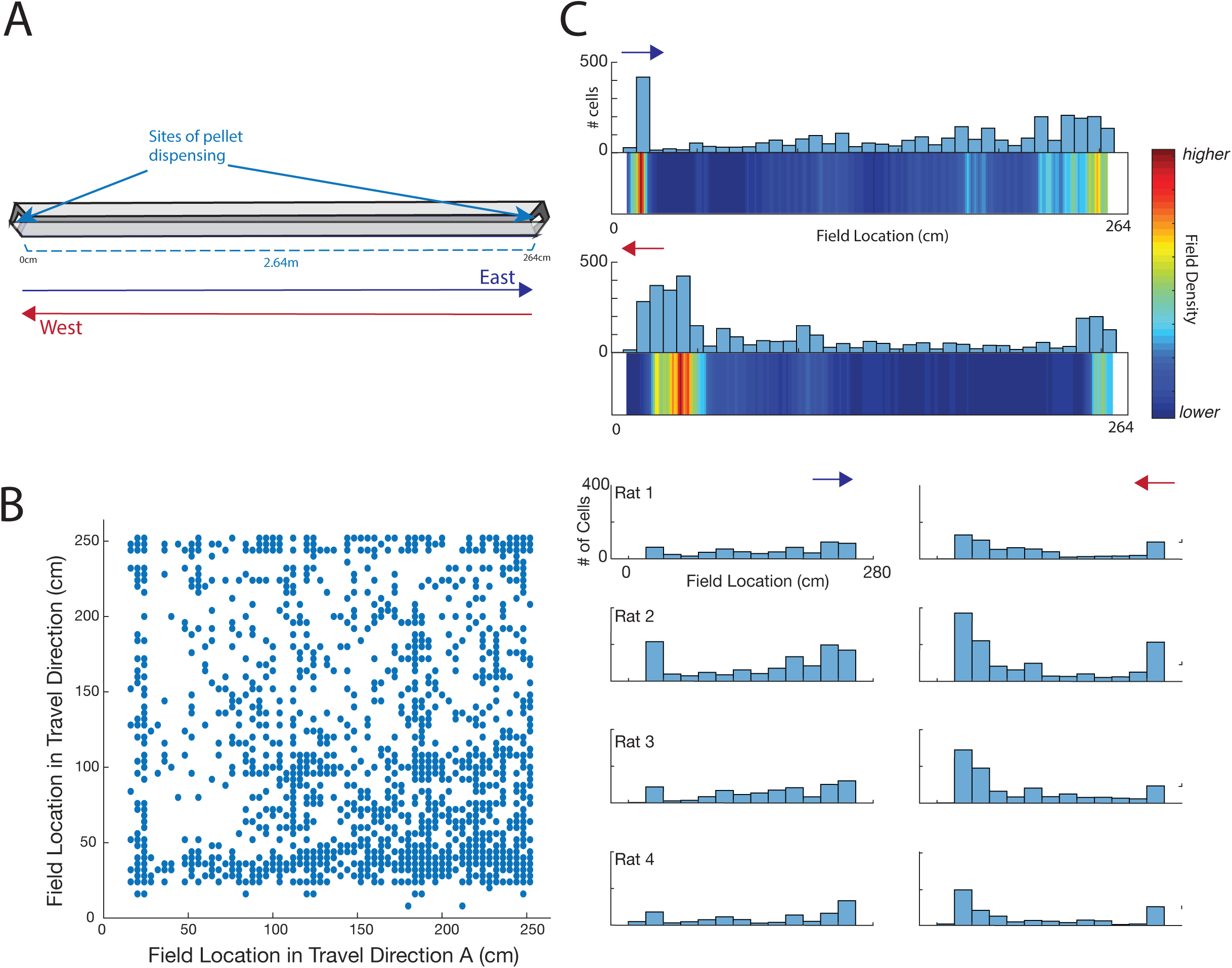
Animals displayed directional place fields while running on a linear track. **a**. Animals were trained to run on a linear track. Animals had to alternate sides to receive a grain pellet reward at either end. **b**. Cells displayed highly directional place fields. Over all cells, there was no correlation between field location in one direction of travel and field location in the other direction (F-test(2415)<.001, p=0.975) **c**. Place field locations were clustered around reward location. Top: Histogram and heat map of place field locations for all sessions and all animals. Top graph indicates one direction of travel and bottom graph the other. Bottom: Place field locations divided by animal number (all sessions per animal)

A cell was considered to be a place cell if the mutual information (MI, see methods) computed in either direction of travel was greater than 95% of MI scores computed 500 times from shuffled cells. Based on these criteria, all trials contained place cells. An average of 77.3+-5% of cells were place cells, with an average of 167.7+-64.6 place cells per session. Our previous work with GCaMP7c in rats identified that the vast majority of expressing cells were principal cells in the stratum pyramidale (Wirtshafter and Disterhoft, 2022).

Consistent with prior literature, we found that recording place cells on a linear track resulted in highly directional place fields (Battaglia et al., 2004; McNaughton et al., 1983; Mehta et al., 1997; Mehta et al., 2000; Muller et al., 1994; Witharana et al., 2016). Of 24 recording sessions, 22 sessions had no correlation (p>0.05) between place field locations in the two directions of travel. Two sessions, from two different animals, had negative correlations between place field locations (F-test(119)=5.89, p<0.05; F-test(100)=13.2, p<0.001). Over all cells, there was no correlation between field location in one direction of travel and field location in the other direction (F-test(2415)<.001, p=0.975) (figure 1B). As expected (Hollup et al., 2001; Lee et al., 2006; Wirtshafter and Wilson, 2020), we found place cells for each direction to be concentrated around reward locations (figure 1c). For all subsequent analysis, we divided place fields by direction of travel.

### Moran’s I can determine spatial clustering

To examine the spatial organization of cells we used a geographic measure of systematic spatial variation, global Moran’s I (GMI), which estimates clustering of like values in a one or two-dimensional space (Cliff and Ord, 1981; Moran, 1950). GMI can provide a quantitative measure of overall, average patterning seen in data. In situations of perfect dispersal, such as a checkerboard pattern, GMI is equal to -1. In situations of perfect clustering, such as a half white and half black board, GMI is equal to +1. In situations of random dispersal, GMI is equal to -1/(N-1), where N is the number of points; thus, the chance value of GMI is negative but as the number of points approaches infinity GMI approaches zero (Figure 2A). It should be noted that while GMI has similarities to the r value from a Pearson correlations in that it ranges from -1 to 1, GMI does not measure the amount of variance that can be accounted for by clustering and is not proportional to any measured variability in the sample. Additionally, seemingly small values of GMI can be both functionally meaningful and statistically significant: because GMI is an average measuring of clustering throughout all points, small areas of tight clustering surrounded by random dispersal may result in a low but highly significant GMI.

**Figure 2:**
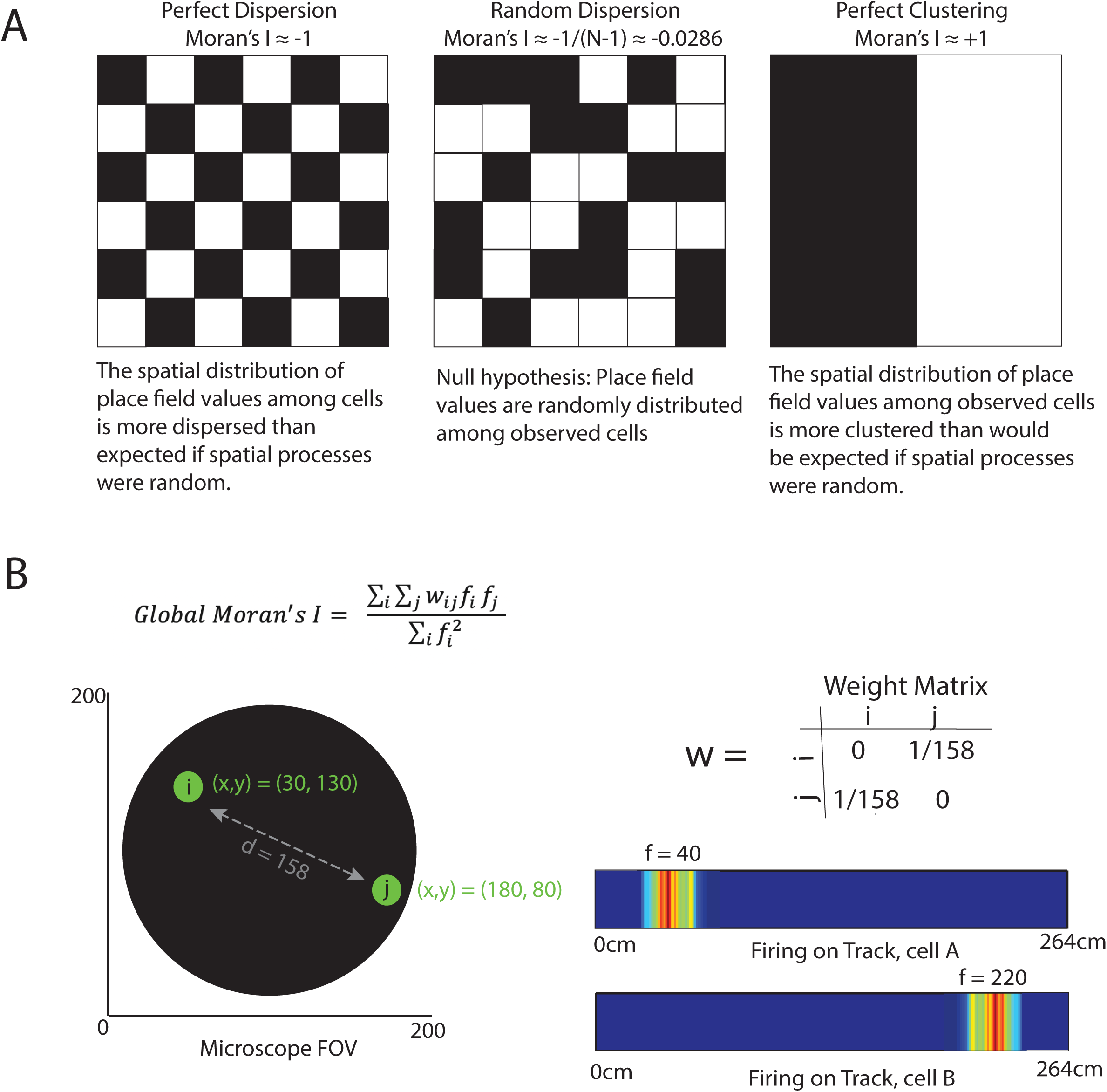
Global Moran’s I can determine spatial clustering. **a**. Examples of Moran’s I. Left: In situations of perfect dispersal, such as a checkerboard pattern, global Moran’s I is equal to -1. Right: In situations of perfect clustering, such as a half white and half black board, global Moran’s I is equal to +1. Middle: In situations of random dispersal, global Moran’s I is equal to -1/(N-1), where N is the number of points; as the number of points approaches infinity global Moran’s I approaches zero **b**. An example of the method by which global Moran’s I was calculated for each recording session. Left: In this example, two cells, ‘a’, location at coordinates (30,130) and ‘b’, at (180, 80) are 158 pixels apart. Right top: Our weight matrix (w) was composed of the inverse of the distance between two cells. Diagonal distances are always zero. Right bottom: The value of ‘f’ in the equation indicates the location of place field center, in this example, cell ‘a’ has a value of 40 and cell ‘b’ has a value of 220.

Global Moran’s I is calculated using a weight matrix which assigns weight to each value based on experimenter’s criterion (see methods): for our analysis, we used a simple distance matrix where the importance of each cell’s field value was based on the inverse distance from that cell to another cell (Figure 2B). A proximal cell would be presumed to have a large effect on the value of close by cells, while a cell further away would have less of an impact.

To determine GMI for each session, we assigned each cell an x,y coordinate location in the field of view based on the brightest recorded location. We then assigned each cell location a number (f) based on the linear location of the place field maximum firing. Each cell thus had two sets of data associated with it: the cell location in an (x,y) coordinate, and a number (f), in cm along the track, indicating place field location (Figure 2B).

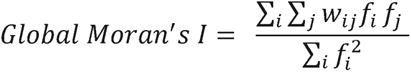

where

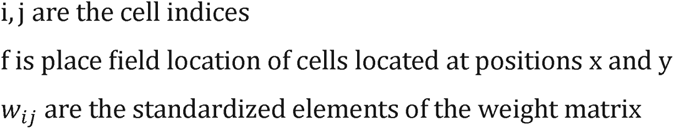

### Place cells are clustered by field location

We first separately examined place fields in each direction of travel, which we will term ‘east’ and ‘west’. We eliminated, from directional analysis, any sessions with fewer than 75 place cells in each direction, leaving 19 sessions for analysis in each direction (at least three sessions from each animal were included). The average number of place fields per session was 140.9+-10.8 going east and 164.3+-11.3 going west. These directional means were not significantly different (t-test T(36)=1.5, p>0.05).

Calculating GMI in the east direction, we found that 18 out of 19 experimental sessions had positive GMI, meaning they were exhibiting clustering, with an average global Moran’s I of 0.0321. The average values of GMI were positive for all animals, with the mean value for each rat being 0.0087, 0.0340, 0.0392, and 0.0436 (Figure 3a- b). Assuming, very conservatively, that GMI should be zero if place fields were randomly distributed (given samples of our size it should actually be below zero; for instance, in a session with 75 place cells, a random global Moran’s I would be -1/n-1 = - 1/74 = -0.14) the odds of 18 or more days having a positive I are p<0.001. To determine whether these values of GMI were significant for individual sessions, we computed GMI 1000 times after shuffling the cell’s place field locations. Importantly, 14/19 sessions had significant GMI values (p<0.05 as compared to the same cell location data with shuffled field locations, bootstrapped 1000 times) (Figure 3c-e). Strikingly, only one session had a GMI value less than the shuffled values (although not significantly less than), and all other 18 sessions had a GMI values above 75% of shuffled values for that session.

**Figure 3:**
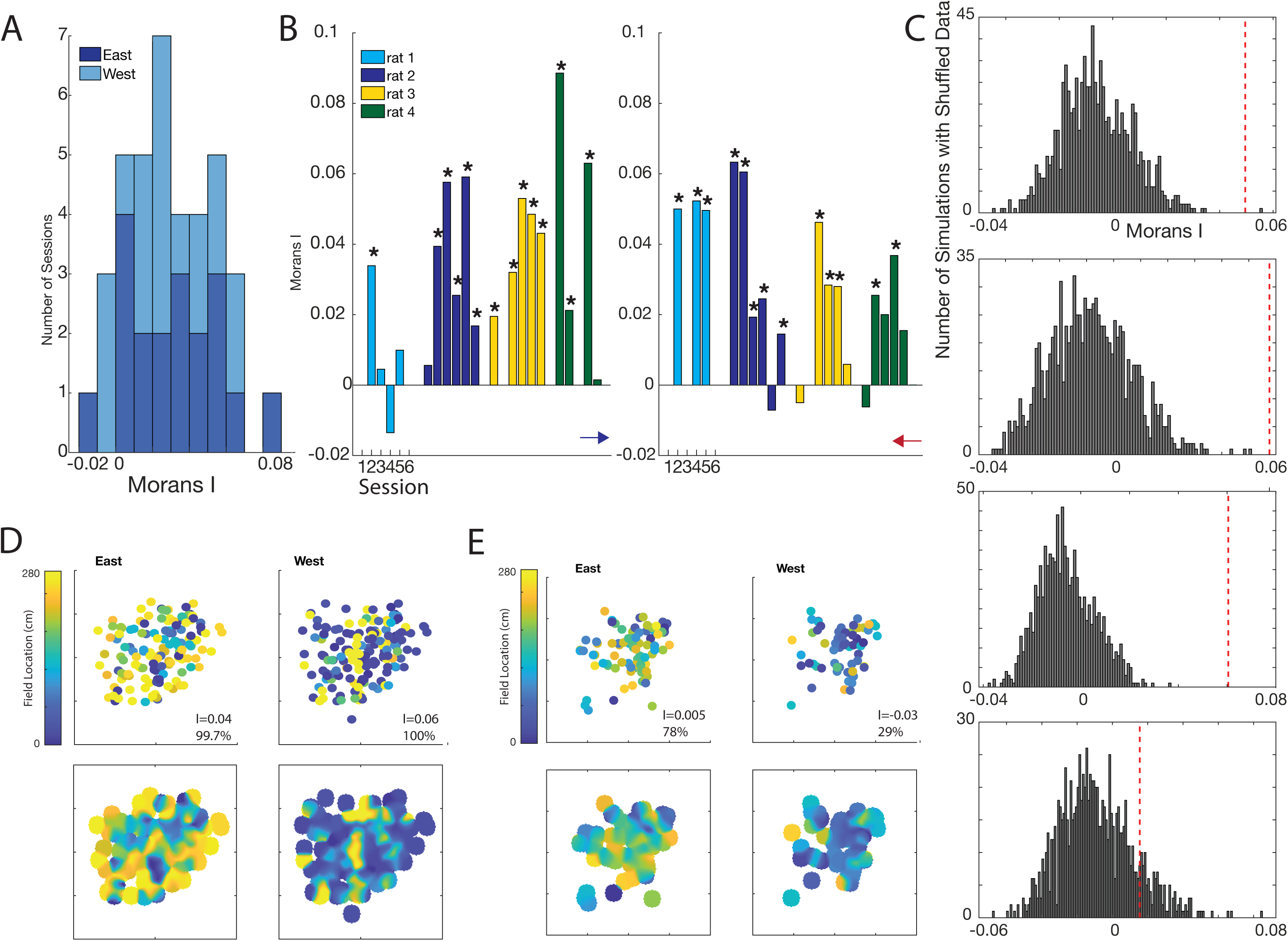
Place cells are clustered by field location. **a**. Stacked bar graph of global Moran’s I values for all sessions. Dark blue represents sessions in the eastern direction and light blue represents sessions in the western direction. 34 out of 38 sessions exhibited positive global Moran’s I values, with 18/19 positive sessions in the east direction and 16/19 positive sessions in the western direction. **b**. Global Moran’s I values for individual sessions of each rat. Left graph is eastern direction and right graph is western direction. Stars indicate the global Moran’s I value is statistically significant (p<0.05) compared to shuffling place field values 1000 times and re-computing GMI. **c**. Four examples of global Moran’s I values compared to those obtained after shuffling place field location 1000 times and re-computing Moran’s I. The red dotted line indicates the actual GMI value for the session. In the top three graphs, GMI was significant compared to shuffled values (p<0.05), while the GMI in the bottom graph was not (p=0.11). **d**. Examples of a session with positive and significant global Moran’s I values. The east and west direction for each cell is shown. The upper plot shows the imaged field of view, and each circle represents an imaged cell. Color of each dot indicates the location of the place field for each cell. On each graph, the GMI value is displayed along with the percent of shuffled and bootstrapped values GMI was greater than. The pictured session had significant GMI values in both directions of travel. Bottom panels are the same as top panels but blurred for easier pattern identification while viewing. Visual examination of the figures shows a visually small but highly statistically significant tendency for cells with similar place fields to be clustered. Note in the left panel, when the animal is traveling east, most cells have place fields approaching the award location (shown in yellow), while, when traveling west, most cells have place fields approaching the award location (shown in blue). Despite this, clustering is seen most clearly in cells with fields proximate to the just-visited reward location: “blue” cells are more clustering on the left, with “yellow” cells more clustered on the right. This phenomenon is discussed in the section “Clustering occurs in patches and clustering is most common in cells with place fields by recently rewarded locations”. **e**. Same as D but for a session which did not have significant GMI values in either direction.

In the west direction, we found that 16 out of 19 experimental days had positive GMI, with an average GMI of 0.0275. This average value for I was not significantly different than the value in the east direction (t-test T(36)=0.59, p>0.05). Individual rats had mean GMI values of 0.0292, 0.0506, 0.0207, and 0.0183 (Figure 3a-b). Similar to travel in the east direction, odds of 16 or more days having a positive I are p<0.005. Of these 19 days, 13 of them had significant GMI values (p<0.05 as compared to the same cell location data with shuffled field locations, bootstrapped 1000 times) (Figure 3c-e).

Again, only one session had a GMI value less than the shuffled value (although not significantly). There was no correlation between the GMI values in a session in the two directions of travel (F(16)=3.7, p>0.05).

As stated, because GMI is an average measuring of clustering throughout all points, small areas of tight clustering surrounded by random dispersal may result in a low but highly significant GMI (as seen in Figure 3c-e). In order to determine local, rather than general, geographic data, it is necessary to use an additional statistic, as presented later.

### Clustering values are not the result of properties/artifacts of calcium imaging

We considered whether the significant GMI values could be the result of mistakenly attributing a single cell’s calcium events to multiple cells, a possibility previously considered by other authors (Dombeck *et al*., 2010). If this occurred, there would appear to be a cluster of cells in one imaging location with the same place field (thereby causing a high global Moran’s I), when these clustered cells were, in fact, a multiple count of a single cell.

We re-computed GMI including only cell relationships that were greater than 0.01mm apart (slightly greater than the radius of an average rat pyramidal cell (Zhuravleva et al., 1997)). This calculation would help alleviate the contribution of tight clusters formed by double counted cells. When we eliminated these connections, 14/19 sessions in the east direction and 14/19 sessions in the west direction had significant GMI values versus shuffled data, compared to 14/19 and 13/19 in the east and west directions, respectively, without removal of these connections. Eight of the measured 39 global Moran’s I values actually increased without inclusion of the connections equal or less than 0.01mm (Figure 4A), and there was no difference in the mean values (0.0298+-0.024 versus 0.0228+-0.019 for no connections removed versus connections <=0.01mm removed, t-test(74)=1.4, p>0.05) of GMI or in the distribution of values (Kolmogorov-Smirnov (KS) test, p>0.05). To take these results a step further, we then eliminated any connections less than 0.02mm, greater than the twice the radius of an average pyramidal cell. Although these re-computed GMI values were significantly different than the original values (t-test and KS-tests, p<0.05), 12/19 eastern sessions and 8/19 western sessions still had significant global Moran’s I values, with 5 sessions actually increasing their GMI (Figure 4A).

**Figure 4:**
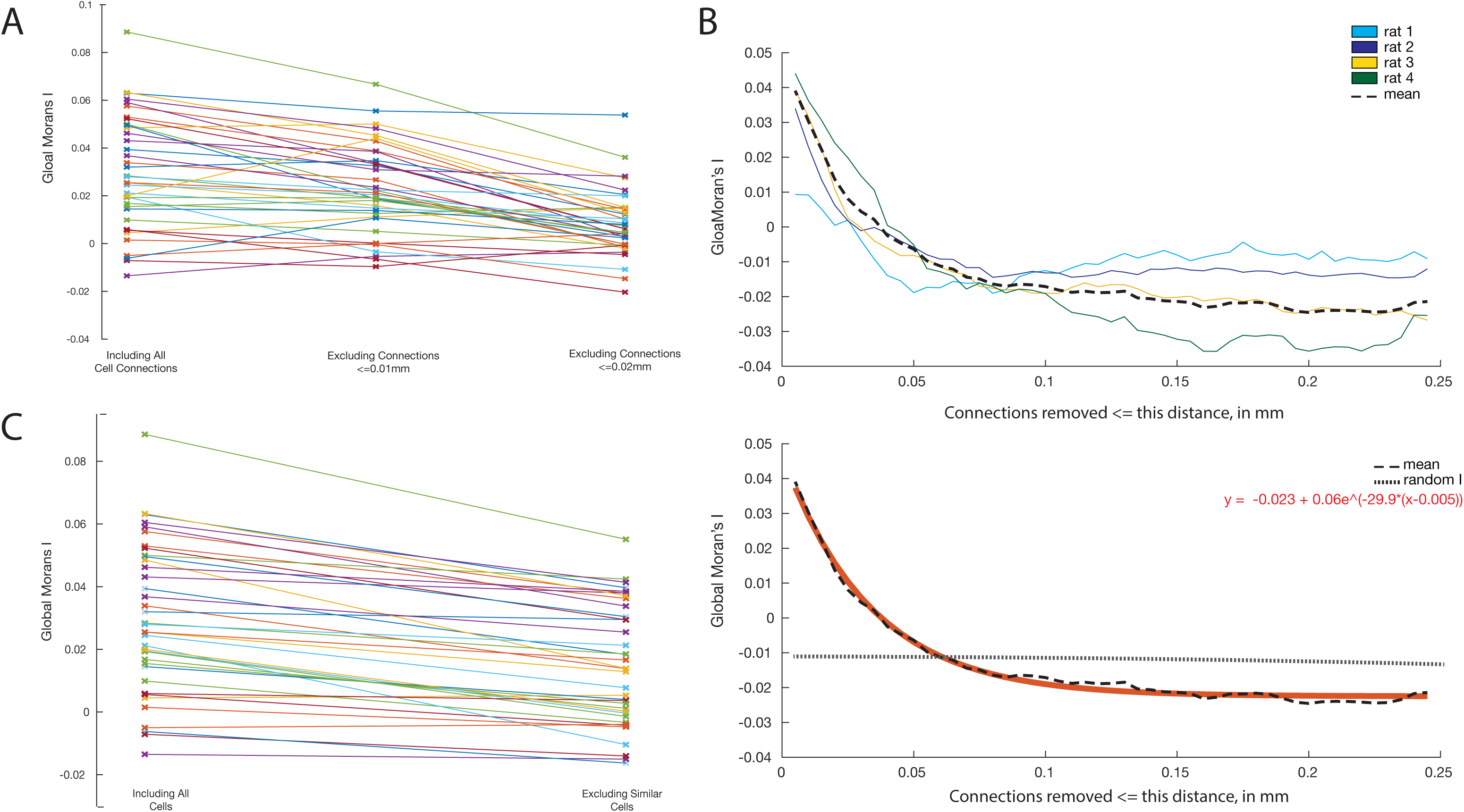
Clustering values are not the result of properties/artifacts of calcium imaging. **a**. Graph showing changes global Moran’s I values from all sessions when eliminating no connections, after eliminated connections of cells <=0.01mm apart, and after eliminating connections <=0.02mm apart. Without removing any connections, 14/19 and 13/19 in the east and west directions, respectively, had significant Moran’s I values. When we re-computed global Moran’s I including only cell relationships that were greater than 0.01mm apart, 14/19 sessions in the east direction and 14/19 sessions in the west direction had significant Moran’s I values, Eight of the measured 39 GMI values actually increased without inclusion of the connections equal or less than 0.01mm, and there was no difference in the mean values (0.0298+-0.024 versus 0.0228+-0.019 for no connections removed versus connections <=0.01mm removed, t-test(74)=1.4, p>0.05) of Moran’s I or in the distribution of values (Kolmogorov-Smirnov (KS) test, p>0.05). We eliminated any connections less than 0.02mm, p<0.05), 12/20 eastern sessions and 8/19 western sessions still had significant GMI values, with 5 sessions actually increasing their GMI. **b**. Graph showing recalculated global Moran’s I while removing connections at less than or equal to varying distances. Top: Displays the graph for of averages from all four animals after removing connections less than or equal to the distance on the x axis. Dotted line shows the mean after averaging all four animals. Bottom: A best fit line of the mean is plotted in red. The value for a chance global Moran’s I given the mean number of points in each session is plotted with a dotted line. We found that only when connections less than or equal to between 0.05mm and 0.055mm—over five times the radius of an average pyramidal cell-- were removed did the average GMI become less than that would be expected by random shuffling. Of note, 8 of the 19 sessions plateaued at Moran’s I values significantly lower than those expected by random shuffling (p<0.05), suggesting that there may be some propensity for cells further apart from each other to have less similar place fields than predicted by random chance. **c**. Graph showing changes in the global Moran’s I value after eliminating all similar cells. Two cells were considered similar if they had both eastern and western place field centers within 10cm of each other, and if the cell centers were within 0.05mm. In each pair of duplicates, we randomly selected one cell to include and one to discard; if a cell was present in more than one pair it was automatically discarded. This led to an average of 10.8+-5.2 cells being discarded per session. We then calculated GMI with these cells removed. Although 36 out of 38 trials saw decreases in GMI, 9/19 sessions in the eastern direction and 10/19 sessions in the western direction still maintained significant (p<0.05) GMI values.

To further validate these results, we calculated GMI while removing connections at less than or equal to varying distances in 0.001mm increments (Figure 4B). We found that only when connections less than or equal to between 0.05mm and 0.055mm—over five times the radius of an average pyramidal cell-- were removed did the average global Moran’s I become equal to or less than would be expected by random shuffling.

Of note, 8 of the 19 sessions plateaued at GMI values significantly lower than those expected by random shuffling (p<0.05), suggesting that there may be some propensity for cells further apart from each other to have less similar place fields than predicted by random chance.

Most stringently, we attempted to detect and remove any possible similar “duplicate” cells by eliminating cells with close place fields and locations. Two cells were considered similar if they had both eastern and western place field centers within 10cm of each other, and if the cell centers were within 0.05mm. Although this distance is quite large, at almost 3x the diameter of an average pyramidal cells, we wanted to make sure we were eliminating any possible duplicates. (We recognize that this method also likely eliminated adjacent cells that were non-duplicates, due to the fact that many cells had similar place fields due to the grouping of fields on either end of the track. These criteria are thus particularly strict in that they may remove non-duplicate cells that are nearby and have similar place fields—exactly the type of cells that would have large contributions to GMI). In each pair of duplicates, we randomly selected one cell to include and one to discard; if a cell was present in more than one pair it was automatically discarded. This led to an average of 10.8+-5.2 cells being discarded per session. We then calculated GMI with these cells removed. Although 36 out of 38 trials saw decreases in GMI (Figure 4C), 9/19 sessions in the eastern direction and 10/19 sessions in the western direction still maintained significant (p<0.05) GMI values versus shuffled data.

### Clustering occurs in patches and clustering is most common in cells with place fields by recently rewarded locations

An additional statistic, termed local Moran’s I (LMI) can be used to determine local features of an environment, as opposed to GMI which averages across the entire feature space (Anselin, 1995). Specifically, LMI can be used to determine specific locations, or cells, that are engaged in clustering or dispersion. In other words, local Moran’s I is the measure of systematic spatial variation of individual points (cells) and is mathematically determined as follows:

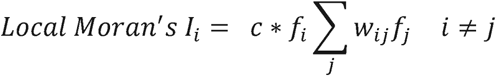

where

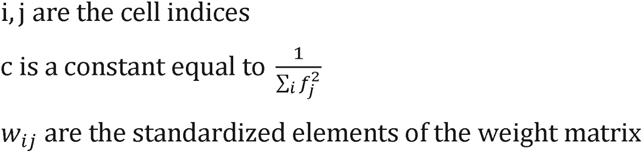

We computed LMI for all cells in both directions. For cells that had place fields in both directions, there was no correlation between the LMI of a cell when the animal was traveling east with the LMI of the same cell when the animal was traveling west (f- test(2567)=0.141, p>0.05) (Figure 5a). There was also no significant difference between local I values in either direction (paired t-test(2568) = 0.27, p>0.05; unpaired t- test(5603) = 0.51, p>0.05, Kolmogorov-Smirnov test=0.025, p>0.05).

**Figure 5:**
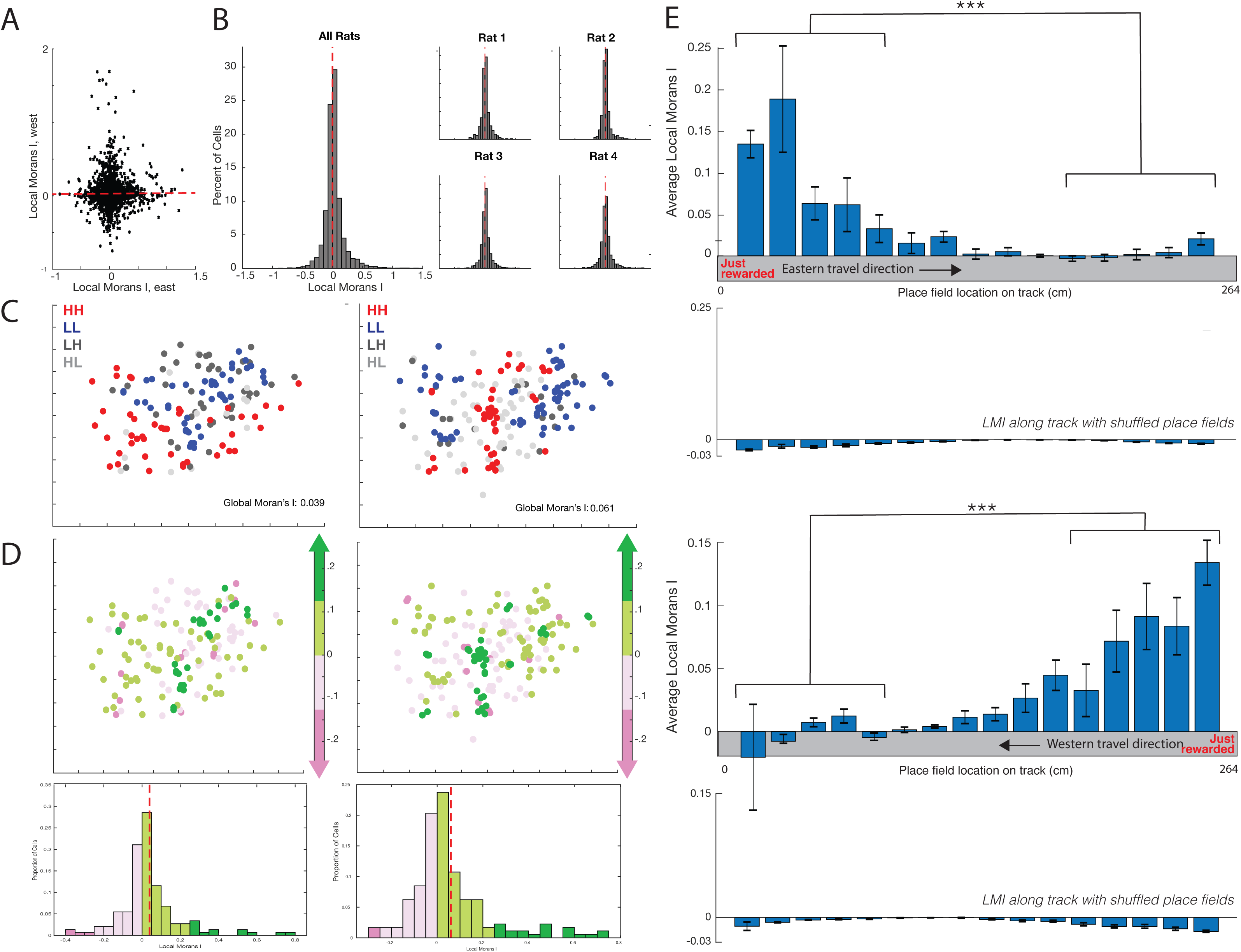
Clustering occurs in semi discrete patches and clustering is most common in cells with place fields by recently rewarded locations. **a**. Scatter plot of local Moran’s I computed for cells when the animal is travelling eastward, versus local Moran’s I computed when the animal is traveling westward. Dotted line is the best fit line with r^2^= -0.0003. For cells that had place fields in both directions, there was no correlation between the local Moran’s I of a cell when the animal was traveling eastern with the local Moran’s I of the same cell when the animal was travelling westward (f-test(2567)=0.141, p>0.05). There was also no significant difference between local I values in either direction (paired t-test(2568) = 0.27, p>0.05; unpaired t-test(5603) = 0.51, p>0.05, Kolmogorov-Smirnov test=0.025, p>0.05). **b**. Histograms showing local Moran’s I values. Red dotted line is at a local Moran’s I value of 0. Axes are the same for all graphs. Left: Histogram of local Moran’s I values for all animals combined. Out of all collected cells, 52.3% had positive local Moran’s I values, meaning their place field location value varied in the direction expected by the values of their neighbors (they exhibited clustering). Overall average local Moran’s I was 0.029+-0.176. Right: Histogram of local Moran’s for individual rats. The rat’s positive rates were 52.8%, 52.7%, 52.5% and 51.5%, with individual rats having average values of 0.027+-.197, 0.030+- 0.166, 0.028+-0.166, and 0.029+-0.196. **c**. Example representation of local Moran’s I from the same session shown in figure 3d: left graph represents cells with place fields when the animal was traveling eastward, right graph is westward. Clustering occurs in semi discrete patches and clustering is most common in cells with place fields by recently rewarded location. The red dots represent cells that have a “high” place field value (on the right side of the track) and would be expected, given clustering, to have a high value based the value of its neighbors (high-high, HH: value is expected to be high and is high). Blue dots represent cells that are expected to have a low place field value (on the left side of the track), and are also expected to have a low value (low-low, LL). High-high and low-low cells are therefore both exhibiting clustering. Dark grey represents cells that have a low value but would be expected to have a high value based on neighbors’ values (low-high, LH), and light grey represents values that have a high value but would expected to have a low value based on neighbors (high-low, HL). Both dark and light grey cells do not display clustering. **d**. An addition representation of local Moran’s I from the same session as in B. Top: Green dots represent cells with positive local Moran’s I values (cells that exhibit clustering—the darker green/the higher the I value, the stronger the clustering based on field location), and pink dots represent cells with negative Moran’s I values (cells that do not exhibit clustering—the darker the pink, the lower the I value, and the more the cells are dispersed based on field location). Bottom: histogram representing local I values from the pictured session. Colors correspond to those on the above graph. Red dotted lines are the mean local Moran’s I value. **e**. Clustering is significantly more prominent in cells with place fields proximate to a just-visited reward location, as can be seen in figure 3d. This figure displays histograms showing local Moran’s I values of cells from all sessions by place field location for travel in the eastern direction (top graph) and western direction (third graph). Error bars represent standard error. Second and fourth graphs are mean local Moran’s I values, in the east and west direction respectively, computed after shuffling cell locations 100 times. In the eastern direction (top graph), the average local Moran’s I was 0.12 for cells with place fields by the recently rewarded location, compared to 0.006 for cells on the opposite end of the track (t-test(1558)=11.35, p<1*10^-28). In the western direction (third graph), the average local Moran’s I was 0.11 for cells with fields by the recently rewarded location, versus -0.001 for cells on the opposite end of the track (t- test(2206)=12.6, p<5*10^-35). Second and fourth graphs: Box plots of local I computed after shuffling cell locations 100 times and recomputing LMI. We compared the distribution of local I values to the distribution that would occur if we shuffled cell locations and recomputed local I values. Shuffled local I values were significantly different in both the eastern and western directions compared to non-shuffled values (Kolmogorov-Smirnov test, p<0.0001 for both directions). Additionally, the mean of local I values for shuffled values was less than 0 for every track segment.

Out of all collected cells in each direction (2657 cells with place fields going east, 3121 going west), 52.3% had positive LMI values, meaning their place field location value varied in the direction expected by the values of their neighbors (i.e., they exhibited clustering). Individual rat’s positive rates were 52.8%, 52.7%, 52.5% and 51.5%. Overall average LMI was 0.029+-0.002, with individual rats having average values of 0.027+-0.007, 0.030+-0.004, 0.028+-0.004, and 0.029+-0.006 (Figure 5b-d).

When cell location was shuffled and local I values were recomputed, only 43% of cells had positive local I values, with a mean local I value of -0.007+-0.002. The shuffled distribution was significantly different than the unshuffled distribution (Kolmogorov-Smirnov test, p<1*10^-14 and unpaired t-test(6473)=7.7 p< 1*10^-13). As seen in Figures 5c-d, clustered cells form small discrete ‘islands’ surrounded by cells that do not participate in clustering.

We then aimed to determine if the LMI of individual cells were correlated to any properties of the cell. We hypothesized that cells with a higher LMI might represent significant locations for the animal. To determine this statistically, we divided the track into three segments: one third of the track by the western reward site, one third of the track between the two reward sites, and a third of the track by the eastern reward site (for visualization in figure 5e, we have further divided the track into 15cm segments). We found that for each animal, the LMI was significant for cells that had place fields by the location the rat was just rewarded at (all t-test, p<0.05). Overall, across all animals, in the eastern direction, the average LMI was 0.12 for cells with place fields by the recently rewarded location, compared to 0.006 for cells on the opposite end of the track (t-test(1558)=11.35, p<1*10^-28). In the western direction, the average LMI was 0.11 for cells with fields by the recently rewarded location, versus -0.001 for cells on the opposite end of the track (t-test(2206)=12.6, p<5*10^-35) (Figure 5e).

We additionally compared this distribution of values to the distribution that would occur if we shuffled cell locations and recomputed local I values. Shuffled LMI values were significantly different from actual values in both the eastern and western directions (Kolmogorov-Smirnov test, p<0.0001 for both directions). Additionally, the mean LMI values for shuffled values was less than 0 for every track segment (Figure 5e).

For travel in the east direction, cells in 15/19 sessions showed a significant correlation (p<0.05) between LMI values and distance from reward location: LMI values were higher for cells that had place fields in located near the just visited reward site. Out of the sessions that had significant global Moran’s I values, 13/14 sessions displayed this correlation. In the western direction cells in 12/19 sessions showed the same significant correlation (p<0.05, with 3 additional sessions with p values between 0.05 and 0.06). Of the sessions with significant global Moran’s I values in the western direction, 12/13 showed a significant correlation (with the 13^th^ having a p value = 0.057). A visual demonstration of this phenomenon can be seen in figure 3d: while cells are more likely to have a place field approaching reward, clustering is more prominent in cells with fields proximate to the just-visited reward location.

### Place field location is most influenced by place fields of neighboring cells within 0.04mm

Using calculations of LMI, we can determine at what distances cell place fields are likely to have a similar field location as nearby cells. To perform this calculation, we can compute LMI while considering or excluding specific relationships between cells. To do this, we can set the weight matrix value for two cells to be 0 if we do not wish to consider that relationship (see methods). We first computed LMI when only including cells that were within an interval 0-0.025mm away, followed by 0.025-0.05mm away, and so on (Figure 6a). We found, as expected, that a cell’s LMI was most likely to be high when calculated including cells closest to it (i.e., cells are more likely to have similar place fields as their close neighbors than to other cells further away). Based on a best fit line (Figure 6a), we observed that the LMI does not fall to chance values until a distance of about 0.12mm. This suggests that cells are no more likely to have similar place fields as cells 0.12mm away than if the place fields were shuffled.

**Figure 6:**
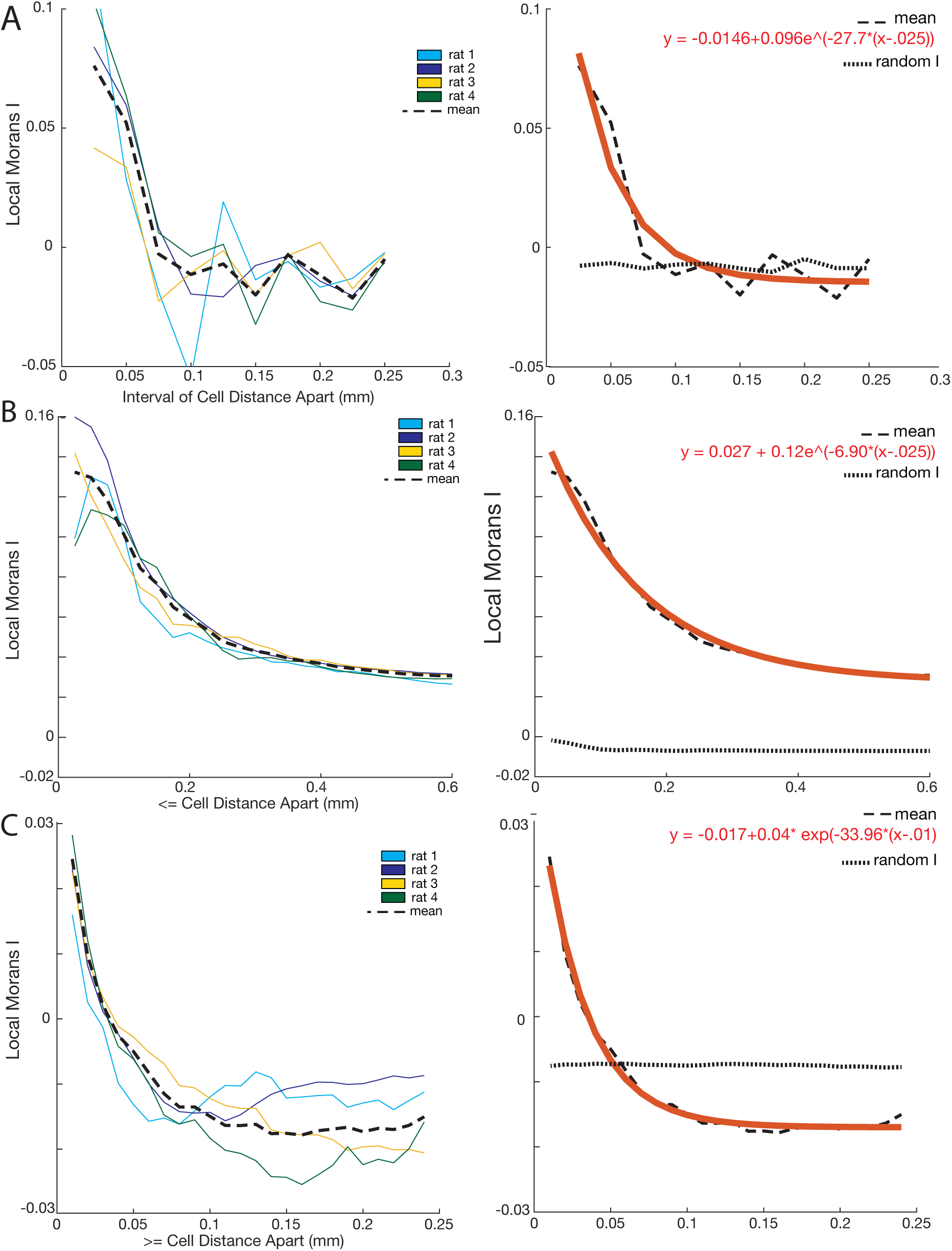
Place field location is most influence by place fields of neighboring cells within 0.04mm. **a.** Local Moran’s I was computed when only cells at particular intervals were included (interval 0-0.025mm away, followed by 0.025-0.05mm away, etc, at 0.025mm increments). Left: Different colors represent the mean values for different rats, and the black line represents the overall mean. A cell’s GMI was most likely to be high when compared with cells closest to it (i.e., cells are more likely to have similar place fields as their close neighbors than to other cells further away). Right: The overall average local Moran’s I (in black dashed line) with a red best fit line. A random LMI value calculated with shuffled cell positions is indicated with a dotted line. Based on a best fit line, local Moran’s I only passes below chance values at about 0.12mm. **b.** Local Moran’s I was computed including cell neighbors at distances less than or equal to the indicated values (<=0.025mm away, <=0.05mm away, <=0.075mm away, and so on). Left: Different colors represent the mean values for different rats, and the black line represents the overall mean. Right: The overall average local Moran’s I (in black dashed line) with a red best fit line. A random LMI value calculated with shuffled cell positions is indicated with a dotted line. As cells approach 0mm apart, they approach a local Moran’s I value of about 0.1642. **c.** Local Moran’s I computed including cell neighbors that were >0.01mm away, >0.02mm away, >0.03mm away, and so on. Left: Different colors represent the mean values for different rats, and the black line represents the overall mean. Right: The overall average local Moran’s I (in black dashed line) with a red best fit line. A random LMI value calculated with shuffled cell positions is indicated with a dotted line. The average local Moran’s I remained above 0 when excluding cells <0.04mm apart, consistent with the above observation (Figure 5b), again indicating that cells are influenced to cluster by neighboring cells up to about 0.04mm away. The calculated value falls below a random LMI at greater than 0.05mm.

We then aimed to find the maximum LMI that could be expected as the distance between cells approached 0mm. To evaluate this, we computed the LMI while only including cells <=0.025mm away, <=0.05mm away, <=0.075mm away, and so on (Figure 6b). The resulting curve closely fits an exponential, f(x) = 0.027 + 0.115*e*^-6.9028(*x*-0.025)^. This suggests that, as cells approach 0mm apart, they approach a limit of a LMI value of 0.1642.

Finally, we computed LMI including cells that were >0.01mm away, >0.02mm away, >0.03mm away, and so on (Figure 6c). We found that the average local Moran’s I remained above 0 when excluding cells <0.04mm apart, indicating that cells are influenced to cluster by neighboring cells up to about 0.04mm away.

### Distance between cells does not, itself, dictate place field location or cell firing patterns

We then looked to quantify if cell firing patterns were influenced by the cell’s location and place field. Although cells tend to be surrounded by cells with similar place field locations, it is yet unclear whether there is a direct linear relationship between the distance between cells and the distance between their place field locations.

On a large scale (the size of the recording area, about 1.1mm^2^), there was no consistent trend between the distance between cells and the distance between their place fields (r values for the four rats were 0.01, 0.01, 0.04, and -0.09) (Figure 7a). We next wished to determine if there was a relationship between cell proximity and field proximity at close distances. When examining only cells <=0.05mm apart, there was also no consistent relationship between distance between cells and distance between fields (r values of -0.07, 0.17, -0.02, and 0.11 respectively), although one rat (with r=.17) did have a significant correlation (F-test(262)=7.47, p<.01) (Figure 7b). (We also looked at individual sessions directionally—out of 19 sessions, each analyzed in both directions, only 3/38 sessions had a significant positive correlation between cell distance and field distance). Based on findings in previous work (Dombeck *et al*., 2010), we also looked at cells <=0.035mm apart and found no significant correlations between distance between cells and distance between their fields (F-test, all p>0.05).

**Figure 7:**
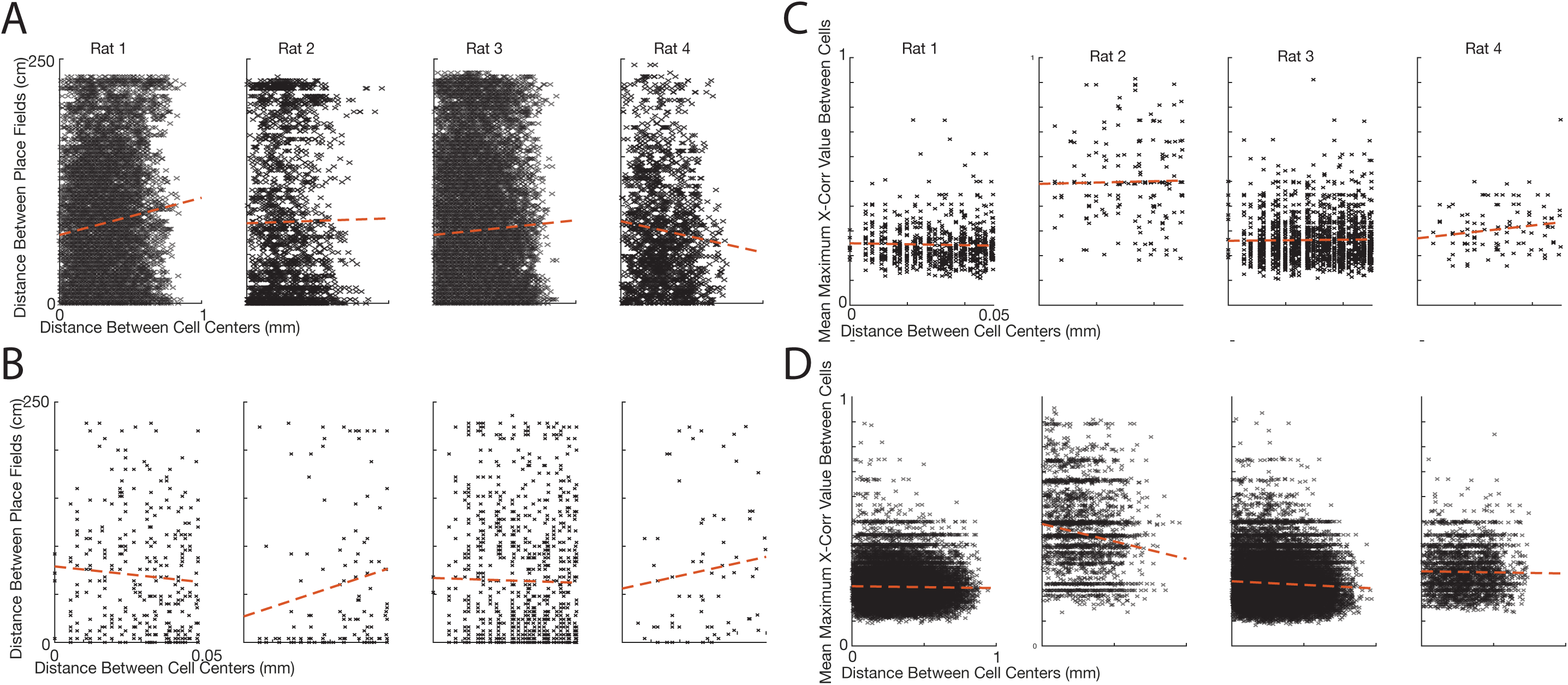
Distance between cells does not, itself, dictate place field location or cell firing patterns. **a**. Linear correlations of the distance between a pair of cell centers and the distance between a pair of place fields for all four rats. Points are vertically jittered for display purposes. Best fit lines are displayed in red. On a large scale (the size of the recording area, about 1.1mm), there was no consistent trend between the distance between cells and the distance between their place fields (r values for the four rats were 0.01, 0.01, 0.04, and -0.09). **b**. Same as A but only computed for cell neighbors <=0.05mm apart, there was also no consistent relationship between distance between cells and distance between fields when cells were <=0.05mm apart (r values of -.07, .17, -0.02, and 0.11 respectively), although one rat (with r=.17) did have a significant correlation (F- test(262)=7.47, p<.01) **c**. Correlations of the distance between a pair of cell centers and their the maximum cross correlation value from the firing of the cell pairs for cell pairs <=0.05mm apart. Best fit lines are displayed in red. There was no consistent relationship between cell distance and cell event patters at <=0.05mm (r values for each rat of -.03, .02., 0.2, and 0.17 respectively). **d**. Correlations of the distance between a pair of cell centers and their the maximum cross correlation over the length of the recording area (about 1.1mm). There was a slight but significant relationship for three animals between cell event correlation and the distance between cells (r values -0.02, -0.14, -0.06, -0.02, all F-test, first three values p<0.0001, last value p>0.05, mean=-0.06)

We then aimed to see if relative cell location had an influence on firing patterns. For each traversal across the track, if a cell had at least three calcium events, we cross-correlated it with all other cells that had at least three calcium events. We then averaged the maximum cross-correlation values, irrespective of lag, for the cell pairs across all traversals. We calculated if there was a significant relationship between maximum cross-correlation values and distance between cells.

Similar to the above analysis, we found no consistent relationship between cell distance and cell event patterns at <=0.05mm (r values for each rat of -.03, .02., 0.2, and 0.17 respectively) (Figure 7c). However, over the entire recording area (about 1.1mm in diameter), we did see a slight but significant negative relationship between cell event correlation and the distance between cells (r values -0.02, -0.14, -0.06, -0.02, all F-test, first three values p<0.0001, last value p>0.05, mean=-0.06) (Figure 7d). The average strength of this relationship did not increase if we excluded cells 0.05mm apart (r values of -0.01, -0.12, -0.05, -0.009, all F-test, first three values p<0.0001, last value p>0.05, mean -0.05), or 0.035mm apart (r values of -0.02, -0.12, -0.06, -0.02, first three values p<0.0001, last value p>0.05, mean=-0.05).

## Discussion

In this study, we use measurements of global Moran’s I and local Moran’s I to demonstrate, for the first time, that place cells in the CA1 of rat hippocampus form “islands” of small clusters of cells with proximate place fields. An individual cell’s place field is most similar to, at the local level, neighboring cells <=0.04mm away, although there is not a strict linear correlation between distance between cells and distance between their place fields. Global patterns of clustering are maintained when examining cells up to ∼0.055mm apart from each other, and cell groups at larger range distances (∼0.25mm) have place fields that tend to be more different than would be expected by chance. We also show, for the first time, that cells with place fields at just-rewarded locations are the cells that predominately form clusters.

While previous work suggested that cells are more likely to have place fields similar to the fields of their neighbors (Dombeck *et al*., 2010; Eichenbaum *et al*., 1989; Hampson *et al*., 1999; Pavlides *et al*., 2019), this is the first study to show that clusters tend to be islands of clustered cells, often comprised of cells with place fields around just-rewarded locations, surrounded by cells not forming clusters. These results also argue against the hypothesis that there is a gradient of field locations across the hippocampus, at least at the scale measured in this study (Eichenbaum *et al*., 1989).

### The unique utility of calcium imaging and Moran’s I

This study, in which we examine the entirety of a 1.1mm^2^ field of view of hippocampus in freely moving animals, has only recently been made possible by the ability to perform calcium imaging in rat hippocampus, in which over 75% of imaged cells have place fields (Wirtshafter and Disterhoft, 2022). Previous attempts to determine the organization of HPC place cells used either extremely local electrophysiology recordings (Eichenbaum *et al*., 1989; Hampson *et al*., 1999; Redish *et al*., 2001), or imaging in the mouse that relied on looking for patterns in a 200x100um field of view with an average of 17 place cells per rerecording (Dombeck *et al*., 2010).

The small area of these recordings, combined with a low number of place cells and the use of correlation and regression analysis, may have obscured the larger organization of HPC cells, as well as local clustering patterns that can be identified with the use of Moran’s I.

The use of Moran’s I to examine the organization of the hippocampus affords several advantages. First, it provides a quantitative measure of both global and local clustering, even when a significant portion of cells do not participate in any clusters. Second, as opposed to using measures of correlation, it allows the examination of spatial organization patterns such as clustering and dispersion that cannot be easily identified using linear relationships, especially in conditions where clusters are not of uniform size and have boundaries that may be difficult to identify. Third, unlike k-nearest neighbor algorithms, it requires no categorization of points, it can operate on continuous variables, and can provide a concrete quantification and measurement of clustering and dispersion. Finally, Moran’s I is a well-established and commonly used measure in geographic and topographic studies, with over 7000 citations on the original manuscript (Moran, 1950) and greater than 24,000 results on Google Scholar.

### Properties of clustered cells make them easily overlooked in previous studies

Issues with other methods, such as regression and nearest neighbor analysis, are compounded when considering that not all cells have positive local Moran’s I values (Figure 5b). Importantly, the cells most likely to have high local I values (Figure 5e-f), and thus to cluster, are cells with place fields proximate to a just-rewarded location. We believe this finding may illuminate some discrepancies between this work and a previous study (Redish *et al*., 2001) that did not find any organization within the hippocampus. In this work, rats ran on either a rectangular track where they were rewarded at 2 out of 4 corners, or on a linear track where they were rewarded at only one end of the track. There was no evidence that place cells recorded on a single electrode bundle (tetrode) were more likely to share place fields with other cells recorded on that tetrode than they were with cells recorded on a different tetrode. Because the animals were rewarded at half the number of locations as in the present study, there was a less frequent post-reward hippocampal response, likely resulting in a minimal amount of clustering compared to the present study.

As explained, the fact that cells divide into small clusters, as opposed to existing in a gradient of field locations, means that correlating the distance between cells with the distance between place fields is not a robust method to determine patterns of spatial organization. We did not find a significant linear relationship between the distance between cells and the distance between their place fields (Figure 7a-b). Despite this, evidence points to cells within ∼0.04mm of each other having more similar place fields than would be expected by chance (Figure 6). (In other words, clustering occurs in a subset of cells within 0.04mm of each other and not all cells participate in clustering. Since a large number of cells do not participate in clustering, we do not see clustering reflected in a correlative analysis that includes all cells.) This 0.04mm measurement is consistent with the previous finding that the distance between cells was related to the distance between fields when cells were <=0.035mm apart (Dombeck *et al*., 2010). However, through the use of local Moran’s I, we showed that this relationship is unique to a subset of cells that participate in clustering: largely cells immediately active after an animal’s award consumption. The limited number of cells that participate in clustering is also likely why, at close distances (<0.05mm), this and other studies have found no or weak correlations between the distance between cells and their firing or calcium event patterns (Figure 7c) (Dombeck *et al*., 2010; Redish *et al*., 2001). The relationship seen across larger (∼1mm) distances is likely not the result of small scale clustering patterns, but rather the result of longer range difference in anatomical inputs (Raisman *et al*., 1966; Risold and Swanson, 1996; Swanson and Cowan, 1977), as well as differences in cell coordination due to the propagation of theta rhythm across the hippocampus (Lubenov and Siapas, 2009).

### Results cannot be explained by artifacts from calcium imaging

Previous studies have suggested the presence of clustering but the authors felt, themselves, that the results were not conclusive because could be the result of artifacts from calcium imaging: specifically, that individual cells’ calcium signals were being double counted due to residual brain or lens motion, or shared neuropil signals (Dombeck *et al*., 2010). We showed that the double counting of cells is not responsible for our results in a number of ways. First, all recordings were corrected for motion artifacts using a stable land mark, such as a blood vessel in the field of view (see methods) (Wirtshafter and Disterhoft, 2022). Second, if any multiple cell centers were identified on the same pixel, we randomly selected only one cell to include in analysis. Although this occurred infrequently, this choice precluded the inclusion of a cell’s events being split between two cells in the exact same location, which could artificially increase Moran’s I.

Furthermore, we demonstrated that even after excluding cell connections larger than the radius of pyramidal cells (Zhuravleva *et al*., 1997) before global Moran’s I computations, a majority of sessions still had significant global Moran’s I values (Figure 4a), and that global Moran’s I does not drop to chance until cell connections less than about 0.06mm are excluded from analyses (Figure 4b). We additionally recalculated global Moran’s I after removing nearby cells with similar place fields. This was a particularly stringent criterion because it, by definition, causes lower global Moran’s I values, but should account for double counting of cells due to both brain/lens movement or shared neuropil signals. Even after this exclusion, a majority of sessions still maintained significant global Moran’s I values (Figure 4c). These results, together, show that it is unlikely that movement or neuropil signals are causing significant global Moran’s I values.

Additionally, if the double counting of cell signals was driving high Moran’s I values, we would expect the same cells to have high local Moran’s I values throughout the entirety of a recording session. However, there was no correlation between a cell’s local Moran’s I value during eastern and western travel on the track (Figure 5a). Additionally, if cell signals were randomly being counted more than once, we would expect high local Moran’s I values to be randomly distributed along the track. However, high local Moran’s I values are very skewed towards one side of the track for both directions of travel and are significantly different than a random distribution (Figure 5e).

We also considered that out-of-plane signals (such as dendritic input) may contribute to the clustering effect, but we believe any effect would be minimal. For out-of-plane interference to influence clustering, it would require that we categorized dendritic signals as calcium events. We believe we have done this, at most, infrequently, for the following reasons: First, our analysis pipeline requires a certain amount of circularity to include a calcium event in analysis, which would preclude the inclusion of long cell processes (see Methods). Second, for dendritic signals to artificially increase clustering, these signals would not only have to be classified as cells, but as place cells with place fields. Given that clustering is most exhibited in cells with place fields after reward locations (Figure 5e), dendritic signals would also have to be primarily classified as place cells with fields after the reward location. If this were the case, we would expect to see an uncharacteristic number of “cells” with place fields after a rewarded location, with the place field count gradually diminishing as the animal moves away from the reward site (as seen for LMI values in Figure 5e). However, our results are consistent with previous studies of animals on rewarded tracks that saw the highest number of place fields when approaching the rewarded location, rather that leaving it, with no evidence of ramping down after visiting a rewarded location (Figure 1c)(Dupret et al., 2010; Hollup *et al*., 2001; Kobayashi et al., 1997; Lee *et al*., 2006; Wirtshafter and Wilson, 2020). This lends further evidence that although we cannot disregard the influence of dendritic input to the clustering effect, any contribution is likely minimal. Similarly, while we cannot eliminate the possibility that interneurons have contributed to this clustering effect, we believe their influence would also be minimal for the above reasons. Furthermore, our previous work indicates that a small proportion of cells labeled with GCaMP7c are interneurons (Wirtshafter and Disterhoft, 2022).

In their entirety, these results provide compelling evidence that artifacts of calcium imaging are not responsible for clustering represented by high Moran’s I values. While concerns regarding residual brain movement and shared neuropil signals have been raised previously in regards to examining cellular organization using calcium imaging (Dombeck *et al*., 2010), this study is the first to show that calcium imaging artifacts are not responsible for clustering results. These findings, combined with the much larger population of place cells than analyzed in previous mouse calcium imaging experiments and the advantages over electrophysiology in determining relative positions of cells, show definitively that clustering occurs in the CA1.

### Cell clustering after reward receipt may reflect increased place cell excitation via dopaminergic inputs

We showed that groups of clustered cells do not have fields randomly distributed in the environment, but rather cells that participate in clustering tend to be place cells proximate to a just-rewarded location. Although further studies will be required to determine the exact behaviors that correlate with this pattern of clustering, there are several possible and speculative circuit and cellular level explanations.

Although a variety of speculations could be made, one interesting set of possibilities involves the role of dopamine in spatial navigation and reward receipt. First, when an animal reaches a goal location, receipt of reward may fuel an increase in hippocampal dopamine release at the specific locations, such as from the brainstem VTA (Kaufman et al., 2020; Kempadoo et al., 2016; Retailleau and Morris, 2018; Sosa and Giocomo, 2021). Because there is sparse dopaminergic innervation of the CA1, these inputs may form small anatomic islands of dopamine release (Kempadoo *et al*., 2016; McNamara et al., 2014; Takeuchi et al., 2016; Tsetsenis et al., 2021). Dopamine release may then result in increased excitability of nearby cells (Lezcano and Bergson, 2002; Zhang et al., 2009). Because place cells are formed from cells with lower spike thresholds (Edelmann and Lessmann, 2013; Epsztein et al., 2011; Harvey et al., 2009; Oh et al., 2003; Valero et al., 2022), place cells for the area of the track visited after reward receipt may therefore be selectively formed from cells in local islands receiving dopaminergic inputs . The effected neurons would slowly lose excitability as the animal retreats from the goal location, explaining why the effect of clustering becomes smaller as place fields move away from the goal location (Figure 5e-f).

Similarly, reverse repay of place cell sequences tends to occur at goal locations after receipt of reward (Ambrose et al., 2016; Bhattarai et al., 2020; Diba and Buzsaki, 2007; Singer and Frank, 2009; Sosa and Giocomo, 2021; Xu et al., 2019). It has been suggested that dopaminergic inputs to the hippocampus play a role in spatial memory formation and consolidation during ripples (Mamad et al., 2017; McNamara *et al*., 2014; Sosa and Giocomo, 2021). Small localized islands of dopaminergic input to the hippocampus, which are ‘activated’ during ripples at reward location, could again serve to create small clusters of cells with place field.

Finally, dopaminergic inputs to the hippocampus are required to reorganize place cells around rewarded locations (Kaufman *et al*., 2020; Mamad *et al*., 2017; Retailleau and Morris, 2018; Sosa and Giocomo, 2021), as well as to mediate spatial learning (Kempadoo *et al*., 2016; McNamara *et al*., 2014; Retailleau and Morris, 2018; Takeuchi *et al*., 2016). It is also known that increased excitation occurs with learning (Oh and Disterhoft, 2020; Oh *et al*., 2003; Whitlock et al., 2006). It is possible that, during learning, an influx of dopamine after reward primes cells in the areas receiving the influx to reorganize around the reward location. Then, even after the task is learned, the subsequent clusters of place cells maintain their field position (Kinsky et al., 2018; Mamad *et al*., 2017; Wirtshafter and Disterhoft, 2022).

In summary of the above ideas, cells with fields in locations immediately after reward receipt tend to be clustered anatomically in areas that are innervated by dopaminergic inputs. Future experiments involving manipulation of reward timing and locations, anatomical studies elucidating exact locations of dopaminergic inputs, and experiments manipulating the functional connectivity between the HPC and dopaminergic areas will be needed to elucidate the exact behavioral, anatomical, and physiological correlates of increased clustering of cells with place fields immediately following reward receipt.

## Conclusion

A challenge in both modern and historic neuroscience has been achieving an understanding of neuron circuits, and determining the computational and organizational principles that underlie these circuits. Deeper understanding of the organization of brain circuits and cell types, including in the hippocampus, is required for advances in behavioral and cognitive neuroscience, as well as for understanding principals governing brain development and evolution (Getting, 1989; Swanson and Lichtman, 2016; Tosches, 2017; Zeng, 2018). Insights into hippocampal organization, as in this study, can elucidate mechanisms underlying motivational behaviors, spatial navigation, and memory formation.

## Acknowledgements

This work was supported by an NIA T32 (T32- AG020506/AG/NIA), an NIA R37 (R37-AG008796/AG/NIA) and an NINDS R01 (R01 NS113804/NS/NINDS). We would like to thank all members of the Disterhoft lab, especially Matthew Oh, Craig Weiss, and Kent Park. We would also like to David Wirtshafter for extensive discussion and editing help.

## Author contributions

Investigation, H.S.W.; Formal Analysis, H.S.W.; Writing – Original Draft, H.S.W.; Writing – Review & Editing, H.S.W., J.F.D..; Supervision, J.F.D.

## STAR METHODS

### LEAD CONTACT AND MATERIALS AVAILABILITY

Questions requests for information should be directed to and will be fulfilled by the Lead Contact, Hannah Wirtshafter. This study did not generate new unique reagents.

### EXPERIMENTAL MODEL AND SUBJECT DETAILS

All procedures were performed within Northwestern Institutional Animal Care and Use Committee and NIH guidelines. Four male Fischer 344 x Brown Norway rats (275g to 325g) were sourced from National Institute on Aging colony at Charles River Laboratories. Different data from the same subjects were previously published in Wirtshafter and Disterhoft, 2021 (Wirtshafter and Disterhoft, 2022). The animals were injected with AAV9-GCaMP7c, implanted with a 2mm GRIN lens and run on a linear maze (Figure 1a). Animals were individually housed in an animal facility with a 12h light dark cycle.

### METHOD DETAILS

#### GCaMP7c Injection and Lens Implantation

GCaMP7c injection and lens implantation were completed as reported in Wirtshafter, Disterhoft 2022 (Wirtshafter and Disterhoft, 2022). In brief: Rats were anesthetized with isoflurane (induction 4%, maintenance 1-2%) and a craniotomy was performed at stereotaxic coordinates Bregma AP -4.00mm, ML 3.00mm. 0.06uL of GCaMP7c (obtained from Viogene Biosciences, packaged AAV9 of pGP-AAV-syn- jGCaMP7c-WPRE, titer 1.73*10^13^GC/mL) was injected over 12 minutes (approximate coordinates Bregma AP -4.00mm, ML 3mm, DV 2.95mm relative to skull), and then th syringe was raised 0.2mm and an additional 0.6ul of GCaMP7 was injected. We repeated this process once more and slightly different coordinates in the craniotomy hole, resulting in 4 total injections.

We then aspirated tissue from the craniotomy site using a vacuum pump and 25 gauge needle. Tissue was aspirated up to and including the horizontal striations of the corpus collosum. A 2mm GRIN lens (obtained from Go!Foton, CLH lens, 2.00mm diameter, 0.448 pitch, working distance 0.30mm, 550nm wavelength) was then inserted into the craniotomy hole and cemented in place using dental acrylic. Animals were given buprenorphine (0.05mg/kg) and 20mL saline, taken off anesthesia, and allowed to recover in a clean cage placed upon a heat pad.

Four to six weeks after surgery, animals were again anesthetized with isoflurane and checked for GCaMP expression. If expression was seen, baseplates were attached using UV-curing epoxy and dental acrylic.

#### Behavioral Training

Behavioral training was completed as reported in Wirtshafter, Disterhoft 2022 (Wirtshafter and Disterhoft, 2022). In brief: Animals were food deprived to 85% body weight and maintained at this weight throughout the experiment. Animals were trained on a linear 2.64m linear track from Med Associates with pellet dispensers at both ends. Animals ran back and forth between the sides of the track and received a grain pellet at alternate ends. Each pellet received was considered a trial.

Six sessions from each animal were selection for analysis. Although each animal was run for >= 7 sessions, an animal would occasionally refuse to run or run slowly. For consistency across analysis, we selected the 6 sessions with the most trials to analyze from each animal. On average, the four animals ran 100.3+-33.1 trials per session, with the individual animals averaging 94.7+-35.3, 85.5+24.0, 114.5+-19.8, and 106.2+-41.2 trials per session.

Because the runs with the most trials were analyzed, trials were not always consecutive. The analyzed sessions were as follows for each animal:

*Animal 1:* Days 1, 2, 3, 7, 8, 9 *(run across 9 days)*

*Animal 2:* Days 1, 2, 6, 7, 9, 13 *(run across 13 days)*

*Animal 3:* Days: 1, 2, 11, 14, 15, 16 *(run across 16 days)*

*Animal 4:* Days: 1, 2, 10, 14, 15, 16 *(run across 16 days)*

#### Calcium Imaging

Calcium imaging was completed as reported in Wirtshafter, Disterhoft 2022 (Wirtshafter and Disterhoft, 2022). Briefly, calcium imaging was done using UCLA V4 Miniscopes (Aharoni and Hoogland, 2019b; Silva, 2017), assembled two 3mm diameter, 6mm FL achromat lens used in the objective module and one 4 mm diameter, 10mm FL achromat lens used in the emission module.

### QUANTIFICATION AND STATISTICAL ANALYSIS

Means are presented as mean+-standard error. Overall averages are computed by finding average per animal then averaging those values. All analysis code is available at https://github.com/hsw28/ca_imaging.

#### Position and speed analysis

Position was sampled by an overhead camera at 30Hz. Position tracking was done post-recording using Bonsai (Lopes et al., 2015). Position on the track was defined as the X coordinate only. Position was then converted from pixels/s to cm/s. Speed was calculated by taking the hypotenuse of the coordinates one before and after the time of interest. Speed was then smoothed using a Gaussian kernel of 1s.

#### Video pre-processing and cell identification

Video pre-processing and cell identification were performed as reported in Wirtshafter, Disterhoft 2022 (Wirtshafter and Disterhoft, 2022). In brief: Videos were recorded with Miniscope software at 15 frames/second. Video processing was done CIATAH software (Corder et al., 2019). Videos were down sampled in space and normalized by subtracting the mean value of each frame from the frame. Each frame was then normalized) using a bandpass FFT filter (70-100cycles/pixel) and motion corrected to a using TurboReg (Thevenaz et al., 1998). Videos were then converted to relative florescence (dF/F_0_); F_0_ was the mean over the entire video.

Cells were automatically identified using CIATAH (Corder *et al*., 2019) using CNMF-E (Zhou et al., 2018). Images were filtered with a gaussian kernel of width 3 pixels and neuron diameter was set at a pixel size of 13. The threshold for merging neurons was set at a calcium trace correlation of 0.65, and neurons were merged if their distances were smaller than 2 pixels and they had highly correlated spatial shapes (correlation>0.8) and small temporal correlations (correlation <0.4).

All cells identified using CNMF-E were then scored as neurons or not by a human scorer. Scoring was also done within CIATAH software in a Matlab GUI. Scoring was done while visualizing and considering a calcium activity trace, average waveform, a montage of the candidate cell’s Ca2+ events, and a maximum projection of all cells on which the candidate cell was highlighted. The relative fluorescence (ΔF/F_0_) local maxima of each identified cell were considered calcium event times. Cell locations were determined based on location of brightest fluorescence.

#### Place cell identification

Place cells were identified using mutual information computed when the animals were running at greater than 12cm/s (Wirtshafter and Wilson, 2020). The mutual information (MI) computed in either direction of travel must be greater than 95% of MI scores computed 500 times from shuffled cells (Kinsky *et al*., 2018). To compute the MI for each cell, the track was divided lengthwise into 4cm bins. The calcium event rate of each cell and the occupancy of the animal were found for each bin. Rate and occupancy were smoothed with a 2cm Gaussian kernel. Mutual information was computed during periods of movement in each direction of travel as follows (Kinsky *et al*., 2018; Olypher et al., 2003; Wirtshafter and Wilson, 2020):

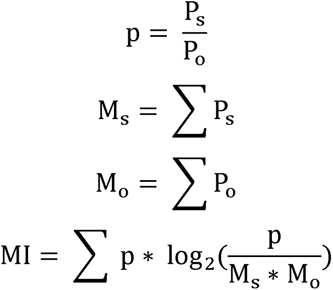

Where:

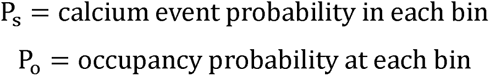

Place field location was determined by the location of maximum firing rate after binning position into 4cm bins.

#### Moran’s I Calculations

Global Moran’s I was calculated using the following formula:

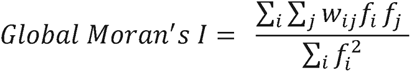

also expressed as:

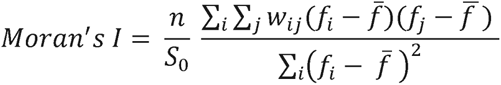

where:

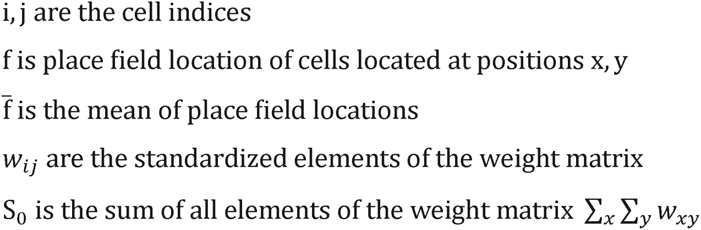

Local Moran’s I was calculated using the following formula:

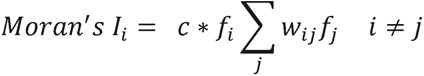

where:

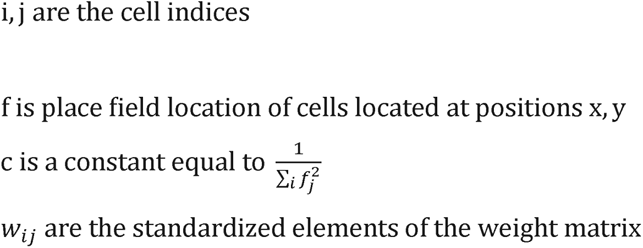

The weight matrices for global and local Moran’s I were calculated the same away: An N by N matrix was created where N = number of place cells recorded in session (C_n_). The matrix was filled in with 1/distance in pixels between C_n_ and C_1:n_, with the diagonal (C_n_ x C_n_) filled in with a value of 0. Because higher values (closer together cells) are weighted more heavily, a value of 0 gives the relationship no weight.

Calculations that required only including cells at a distance greater than, less than, or equal to a certain number of pixels (Figure 4 and Figure 6) were computed as follows: if the distance between the specified cells fell outside the specified values, the distance was replaced with a 0 in the weight matrix. As explained above, a value of 0 gives the relationship between those cells no weight. For figures 6a and 6b, curves were re-computed changing the X variable by 0.025mm per calculation. For figure 6c, the x variable was changed by 0.01mm per calculation. A new weight matrix was created for each calculation.

#### Shuffling and determining significant GMI

To determine which global Moran’s I values were significant, we shuffled the locations of place fields, resulting in a ‘mismatch’ between cell locations and field locations. We then recomputed the weight matrix and Moran’s I. This process was repeated 1000 times. A global Moran’s I value was considered significant if it was greater than 950 (95%) of GMI values computed after shuffling (p<0.05).

For determining random LMI and GMI values for Figure 6 and Figure 4b respectively, curves were recomputed 100x per session after shuffling field locations. Sessions were then averaged together to create the line signifying random LMI or GMI.

#### Cross-correlations

Cross correlations between cell calcium events were performed as follows: sessions were separated into individual trials (one run from one side of the track to the other) of at least 1 second long. Place cells that had 3 or more calcium events during the trial were identified. Thee trial was then divided into 0.01s bins and the calcium event times (‘spike trains’) of each identified cell were cross correlated with each other cell identified in that trial. If the cross correlation for a pair of cells was computed multiple times (they each had >=3 calcium events on multiple trials), the cross-correlation values were averaged together.

### DATA AND CODE AVAILABILITY

All analysis code is public on https://github.com/hsw28/ca_imaging

